# Meta-analytic connectivity modelling of deception-related brain regions

**DOI:** 10.1101/2021.03.09.434551

**Authors:** Sarah K. Meier, Kimberly L. Ray, Juliana C. Mastan, Savannah R. Salvage, Donald A. Robin

## Abstract

Brain-based deception research began only two decades ago and has since included a wide variety of contexts and response modalities for deception paradigms. Investigations of this sort serve to better our neuroscientific and legal knowledge of the ways in which individuals deceive others. To this end, we conducted activation likelihood estimation (ALE) and meta-analytic connectivity modelling (MACM) using BrainMap software to examine 45 task-based fMRI brain activation studies on deception. An activation likelihood estimation comparing activations during deceptive versus honest behavior revealed 7 significant peak activation clusters (bilateral insula, left superior frontal gyrus, bilateral supramarginal gyrus, and bilateral medial frontal gyrus). Meta-analytic connectivity modelling revealed an interconnected network amongst the 7 regions comprising both unidirectional and bidirectional connections. Together with subsequent behavioral and paradigm decoding, these findings implicate the supramarginal gyrus as a key component for the sociocognitive process of deception.

## Introduction

The motivation for researching the complex behavior of deception exists not only to identify mechanisms of sociocognitive functioning, but also to further efforts to detect instances of suspect behavior. Deception is a critical aspect of criminology and forensic/legal decision-making. Deception may be defined as “the act of causing someone to accept as true or valid what is false or invalid” [1]. Deception occurs at various levels of society even becoming apparent in current politics. Specifically, deception occurs in social settings and requires a willful decision from the individual deceiving another [2]. Young, preschool age children are able to comprehend the concept of lying [3], indicating the quotidian nature of deception established early on in cognitive and behavioral development. Psychological assessment of psychopathy even considers one’s ability to lie, deceive, or manipulate [4]. The evolutionary and developmental bases of both verbal and non-verbal deception have previously been reviewed [3]. Moreover, uncovering neural substrates of deception has recently become an important area of research. Brain-based deception research began in attempts to advance traditional polygraph testing [5]. The first report of the neuroanatomical correlates of deception used functional magnetic resonance imaging (fMRI) metrics [6].

In their pioneering publication, Spence et al. [6] had participants answer yes/no questions while undergoing fMRI to investigate the hypothesis that inhibition of truthful responses would be associated with greater ventral prefrontal cortex (PFC) activation. The researchers also investigated if the generation of a lie would be associated with greater dorsolateral PFC (DLPFC) activity. Results showed that lying was associated with increased activation in bilateral ventrolateral PFC (VLPFC) and anterior cingulate cortex (ACC) in addition to medial premotor and inferior parietal cortices.

Langleben et al. [7] utilized the guilty knowledge paradigm to test the hypothesis that participants would activate inhibitory brain regions involved in executive control while withholding a truthful response. Results demonstrated that lying was associated with greater ACC and left parietal cortex activation, replicating Spence et al.’s initial findings [6]. A feigned memory impairment task (where normal individuals pretend to have memory loss) was conducted by Lee et al. [8] showing that malingering was associated with increased activation in bilateral DLPFC, inferior parietal, middle temporal, posterior cingulate cortices, and bilateral caudate nuclei. Further exploration of deception and the brain was conducted by Ganis et al. [9] who investigated well-rehearsed versus spontaneous lies. Both types of lies were associated with greater activation in bilateral anterior PFC and bilateral hippocampal gyri. The aforementioned studies consistently demonstrated converging evidence across differing paradigms that deception involves the prefrontal and anterior cingulate regions of the brain.

As noted, deception has been examined using a wide range of tasks. While there are consistent findings across many studies, some variance exists related to the brain regions involved in deception. It is likely that the neural underpinnings of deception vary based on the act of deception recruiting areas functionally associated with decision making, risk taking, cognitive control, theory of mind, and/or reward processing [10]. Most often reported is activation of prefrontal regions (DLPFC, VLPFC or ventromedial PFC) and ACC, in addition to the inferior frontal gyrus (IFG). Also reported in the literature are the anterior insula, precuneus, inferior parietal lobule (IPL), medial frontal cortex, and regions of the temporal lobe.

Three prior meta-analyses have addressed the issue of variable activation reported during deception. Christ et al. [11] used activation likelihood estimation (ALE) to quantitatively identify regions consistently more active during deceptive responses than truthful responses. ALE pools 3-dimensional coordinates in stereotactic space from task-based brain activation studies. Results identified deception-related activation in the bilateral insula, bilateral IFG, bilateral medial frontal gyrus (MFG), bilateral IPL/supramarginal gyrus (SMG), right thalamus, right ACC, left internal capsule, and left PFC. Further, they found that 10 of 13 peak deception-related regions were associated with working memory, inhibitory control, or task switching, which are all components of executive function.

Lisofsky et al. [12] extended the work of Christ et al. [11] by including “more ecologically valid and interactive experimental paradigms” in their meta-analysis. Lisofsky et al. based their meta-analysis on the idea that deception is both a sociocognitive and executive process, pursing Christ et al.’s [11] finding of deception-related IPL activation that was not correlated with aspects of executive control. Lisofsky et al. [12] found bilateral activations in ACC, IFG, and insula in addition to bilateral activity in IPL, and left MFG. This network was “almost the same network” Christ et al. [11] reported in their work.

The most recent meta-analysis of deception and the brain focused on the distinction between a deliberate attempt to deceive and a true false memory when not telling the truth [13]. Yu et al. [13] also used ALE to separately evaluate deceptive versus truthful responses and false memories versus true memories. Analysis of deceptive versus truthful responses revealed 10 significant clusters primarily in bilateral frontoparietal regions including IFG, superior frontal gyrus (SFG), MFG, insula, SMG, and caudate. The researchers stated that findings discussed in both previous meta-analyses [11,12] were not sufficient to warrant fMRI-use in high stakes legal contexts for detecting deception. They believe their work added the key factor of considering why falsehoods arise (to deceive or not to deceive), not simply if they do.

In the current study, we use the ALE method of coordinate-based meta-analysis [14,15]. By pooling 3-dimensional coordinates, ALE analyzes voxel-wise, univariate effects across the various experiments and generates a probability distribution that is centered at the respective coordinates [16,17]. Building on this meta-analysis, we examine how deception-related brain regions are functionally connected using meta-analytic connectivity modelling (MACM) [15,18-20]. MACM uses regions from ALE to quantify covariance patterns (networks) via patterns of activation reported across a wide range of paradigms [15,18,21]. To our knowledge, this is the first meta-analysis to conduct connectivity analyses in an investigation of deception and the brain. The use of functional connectivity in studies of deception may provide greater insight into its neuropsychological mechanisms, provided that the majority of cognitive processes are supported by various brain networks, rather than single brain regions.

The aims of this meta-analysis are as follows: first, to replicate previously reported brain regions consistently activated during deception across the varying task paradigms; and second, to determine a functionally connected brain network distinct to deceptive behavior versus honest behavior. Our a priori hypotheses are: first, that we would observe activation in prefrontal and memory-related regions of the brain across the various paradigms; and second, that we would observe functional connections involving those regions within the resultant network.

## Methods

### Literature search criteria and study selection

Peer-reviewed articles published prior to August 26^th^, 2020 were selected through searches on PubMed. The Preferred Reporting Items for Systematic Reviews and Meta-Analyses (PRISMA) guidelines [22] were followed, and the selection process is detailed in Fig 1. The initial search keywords used were: (deceptive OR deception OR dishonest) AND (fmri OR magnetic resonance imaging). The following filters were applied to the initial search results on the database: human subjects, adults (18+), and English language. Additional databases (Google Scholar and PsycInfo) were searched via similar terms for articles not on PubMed. Each article was subsequently reviewed (first by abstract, then by full-text) for relevance to the study and inclusion of all following criteria: 1) published between 2005 and 2020, 2) carried out via task-based functional magnetic resonance imaging, 3) at least five healthy (human) adult subjects, 4) peak activations were reported (*x, y, z* coordinates provided in either MNI (Montreal Neurological Institute) space or Talairach; coordinates reported in Talairach space were converted to MNI using GingerALE (version 3.0.2.) [14,17,23], 5) a contrast was reported representing locations of greater activation for deceptive responding as compared to being truthful, 6) contrasts were calculated using a commonly accepted level of significance in a whole brain analysis, and 7) information regarding the task and stimulus material used were reported.

**Fig 1.**
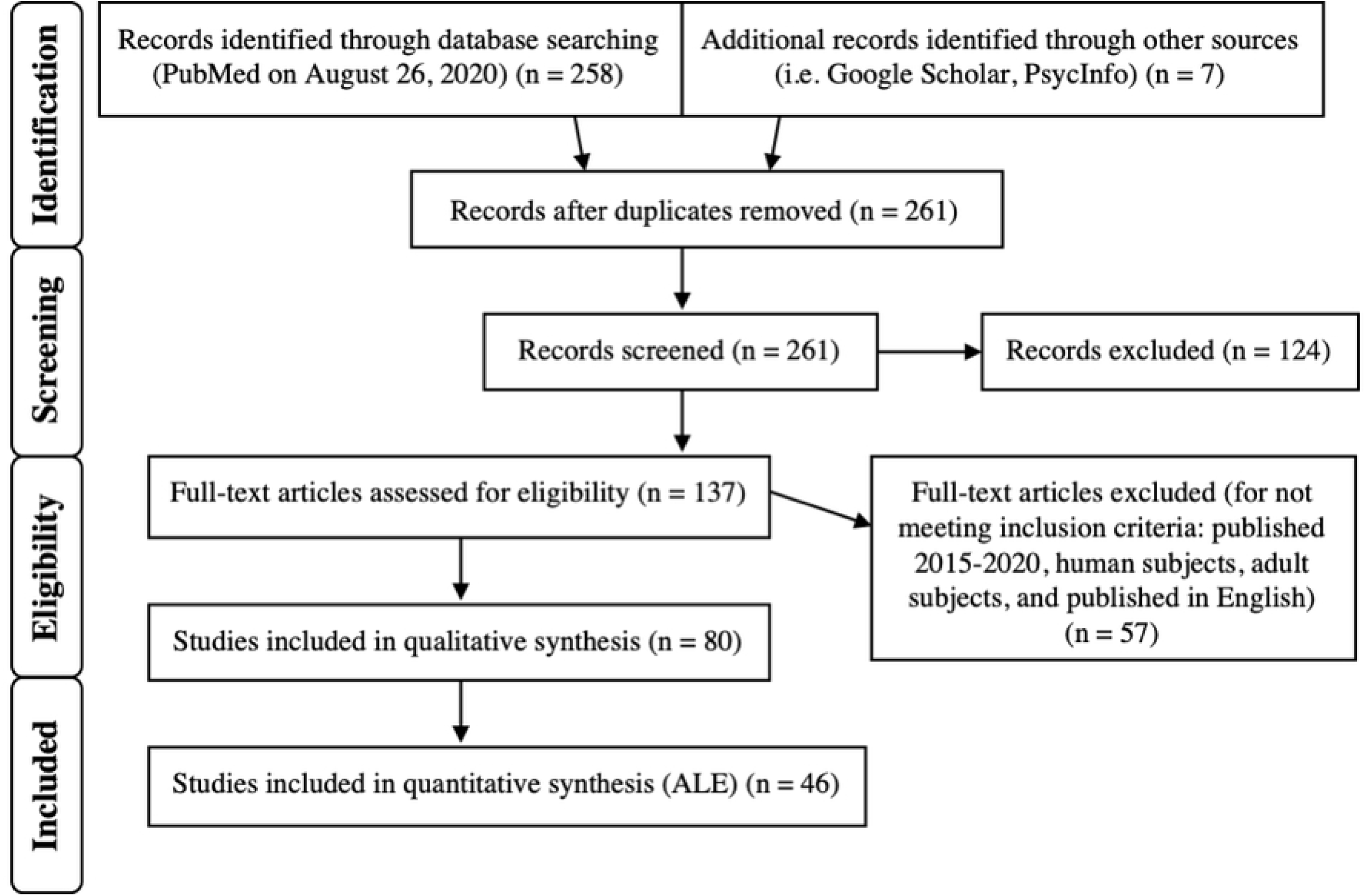
PRISMA Diagram. This diagram depicts the inclusion criteria and study selection process [22].

Any relevant contrast related to deceptive versus honest behavior (D > H) in a relevant article was included to provide a complete analysis of reported contrasts for deceptive or honest behavior. For example, “Lie > Truth” and “Identity Concealment > Control” were both considered comparisons between deceptive behavior and honest behavior. Any article reporting the opposite contrast was included in the supplemental analysis of all contrasts (i.e. “Truth > Lie”). Table 1 details all contrasts included in the D > H ALE and MACM. S1 Table details all contrasts of included articles, including both deceptive > honest and honest > deceptive contrasts.

**Table 1.**
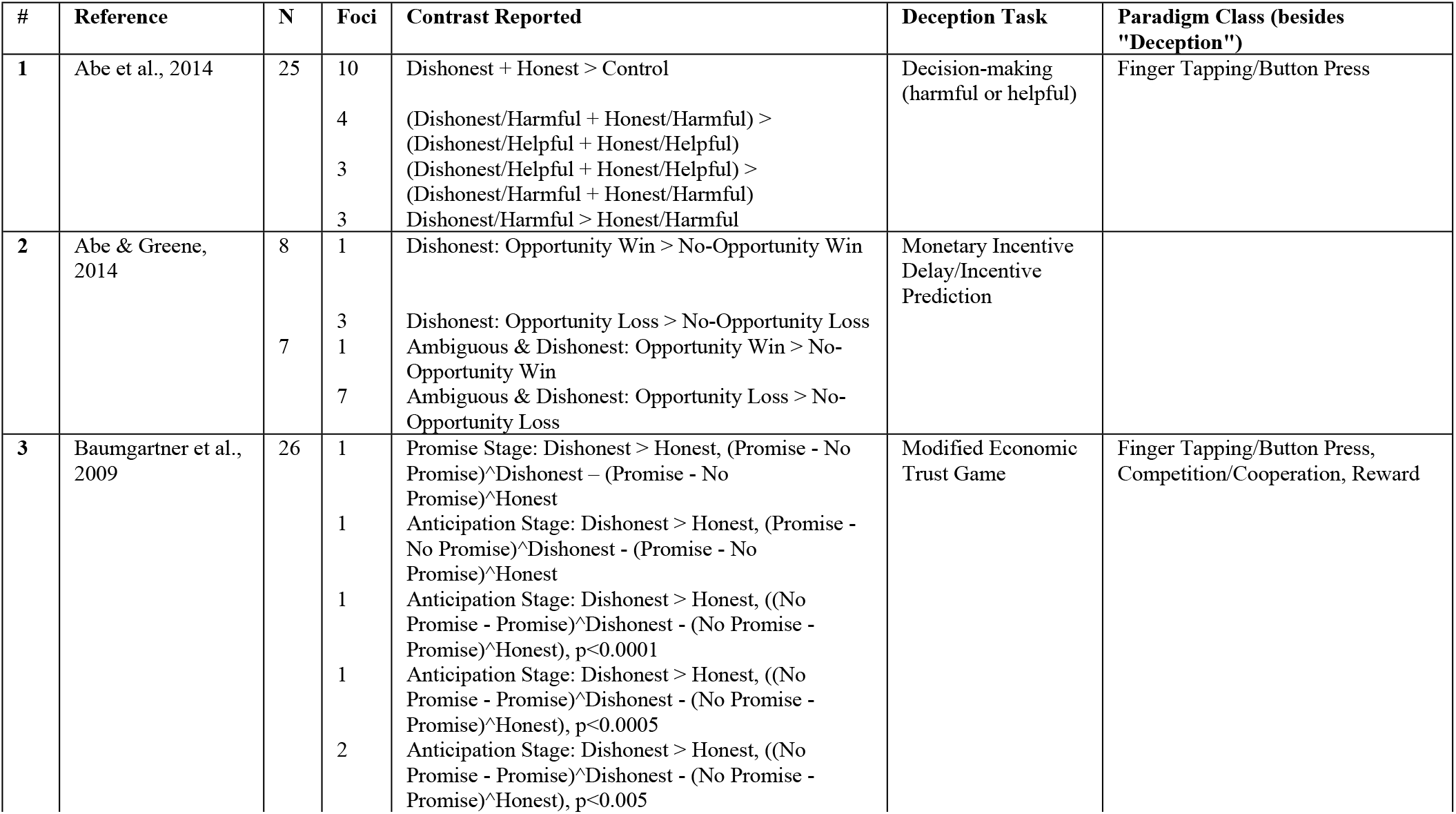
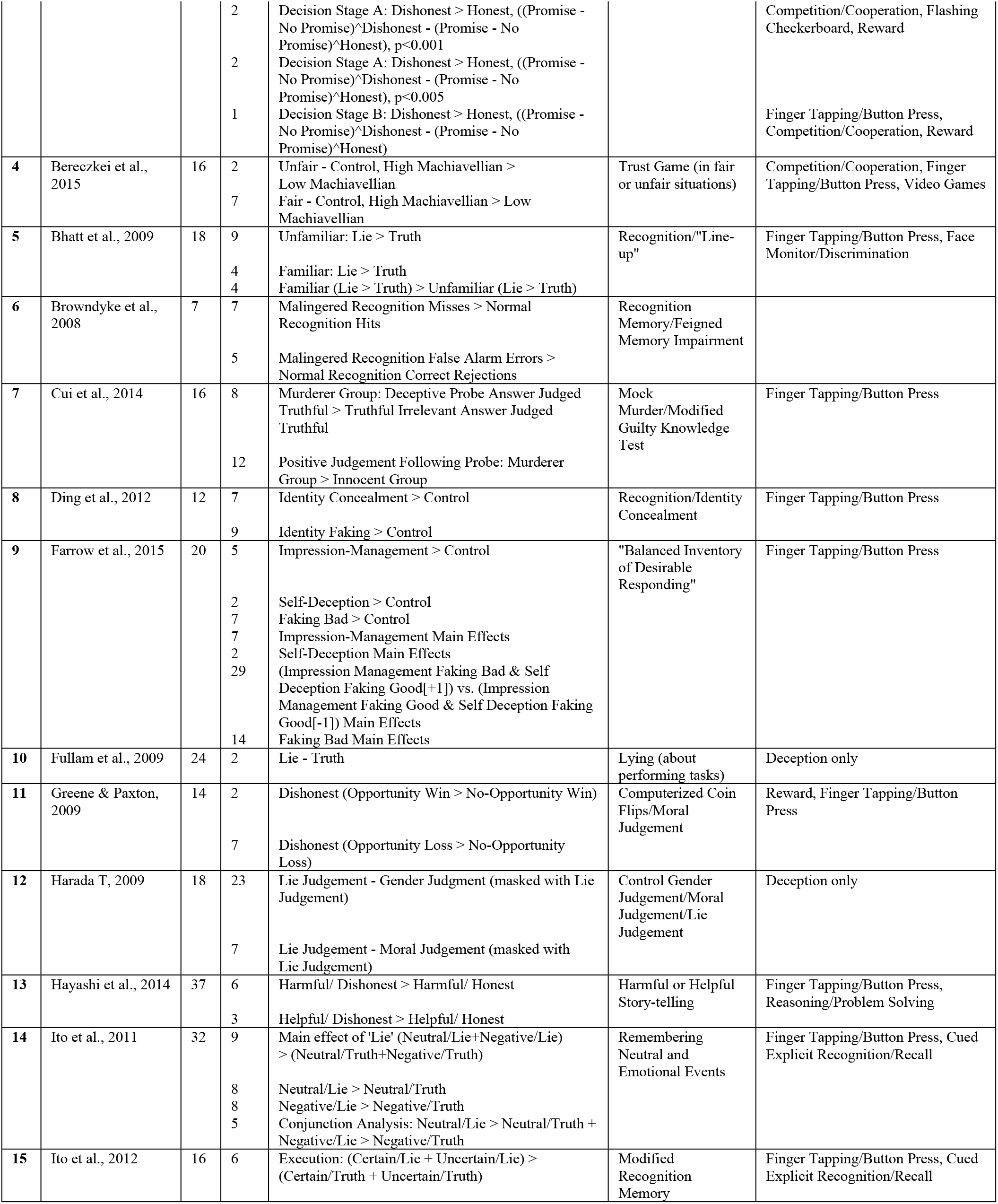
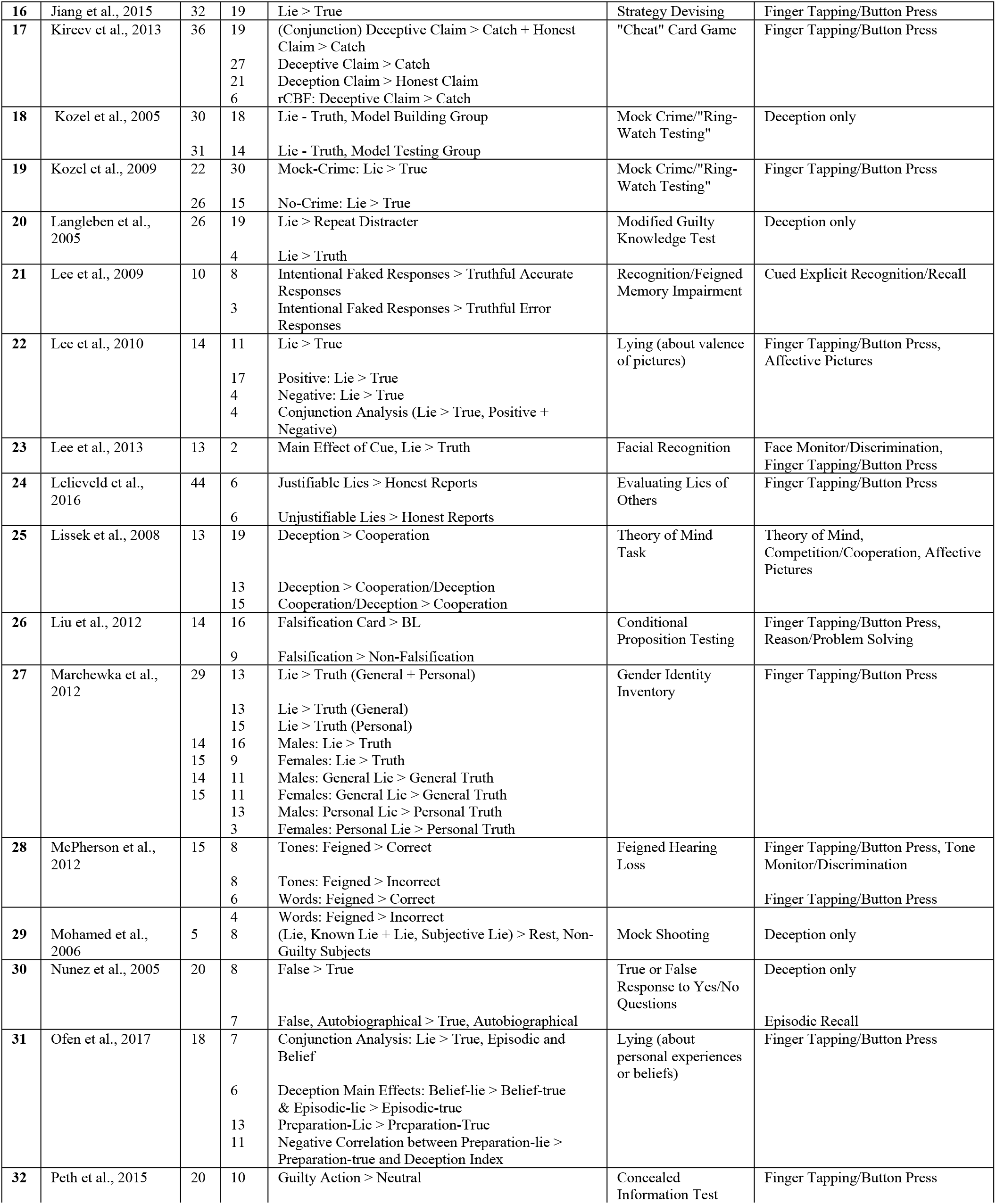
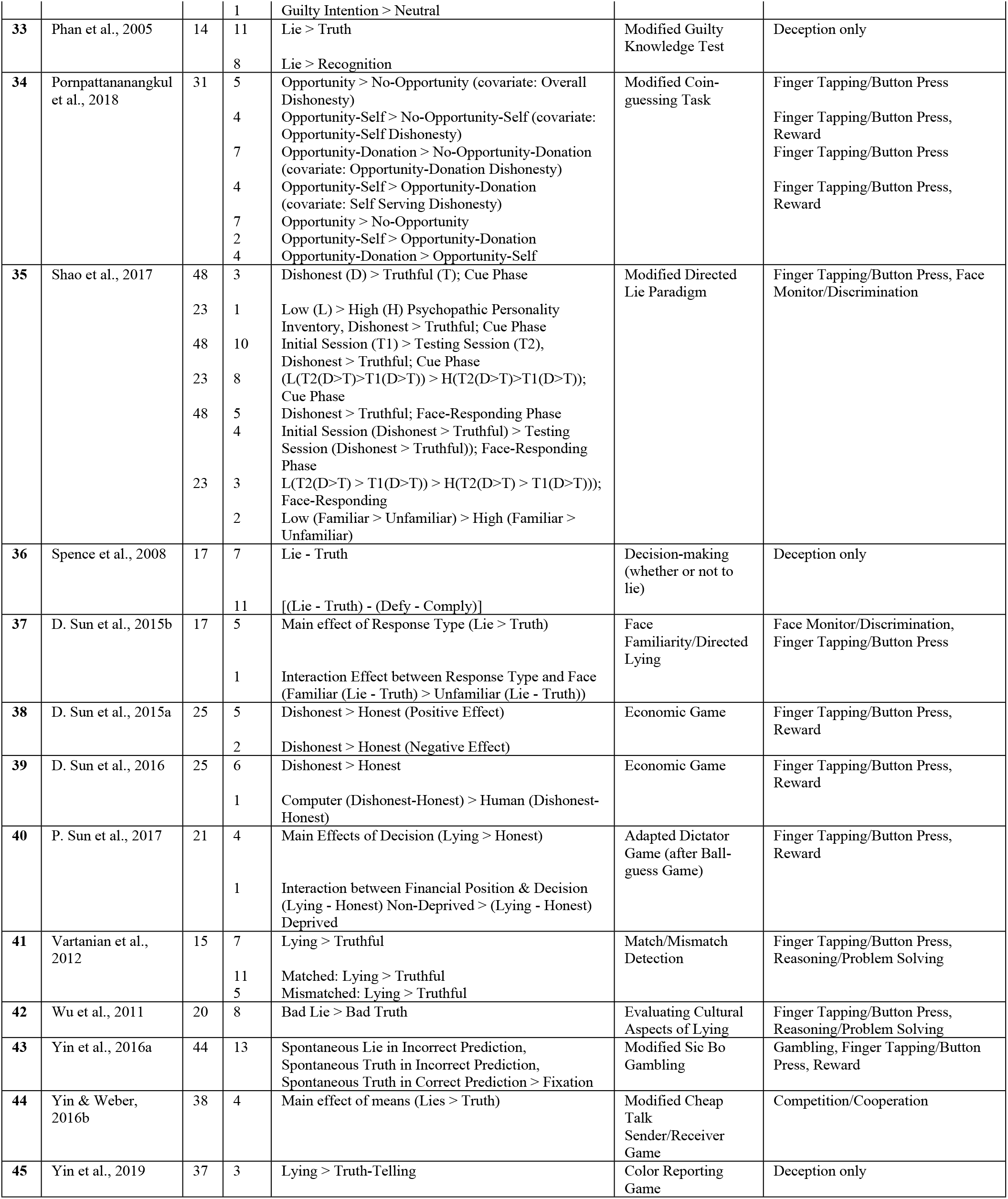
Contrasts included in Deceptive > Honest ALE.

The number of participants, number of reported foci, deception task used, and paradigm class are listed for each reference/contrast.

BrainMap software was used to carry out both ALE and MACM. BrainMap [24] is a database that archives published coordinate-based results in standard brain space from neuroimaging experiments [25]. At the time of analysis, the BrainMap Functional Database contained over 3,400 papers consisting of over 16,900 experiments with over 76,000 subjects and 131,500 coordinate locations. The software used in the following analyses are briefly described here: Scribe (version 3.3) [16,26,27] allows users to submit data and meta-data from selected publications; Sleuth (version 3.0.4.) [16,26,27] allows users to search for and retrieve coordinate data and meta-data from various publications archived in BrainMap; GingerALE (version 3.0.2.) [14,17,23] allows users to carry out ALE-based meta-analyses.

### Activation likelihood estimation

ALE [14,15] was carried out using activation coordinates from the included studies (Table 1) and BrainMap’s GingerALE software (version 3.0.2.) [14,17,23]. The primary ALE conducted and reported was based on deceptive versus honest (D > H) behavior. This D > H ALE included 45 studies and 127 experiments with 977 foci from 2,836 subjects. Subsequent ALE analyses are reported in Supplementary Material (see S2 Table; all contrasts: deceptive versus honest, honest versus deceptive, etc.).

We followed standardized procedures for performing ALE using BrainMap’s software as reported in the GingerALE user manual (Research Imaging Institute, 2013, *http://www.brainmap.org/ale/manual.pdf*). For ALE meta-analysis, a set of coordinates, in addition to any experimental meta-data (identified as suitable for the specific research question), are retrieved via Sleuth. These coordinates are input to GingerALE and smoothed with a Gaussian distribution to accommodate the associated spatial uncertainty (using an estimation of the intersubject and interstudy variability typically observed in neuroimaging studies) [25]. A statistical parameter (the ALE value) is computed which estimates convergence across brain images and measures the likelihood of activation at each voxel in the brain. Additionally, the ALE algorithm calculates the above-change clustering between experiments (random-effects analysis) rather than between foci (fixed-effects) [25]. The ALE value is generated for each voxel and converted into p values for identification of areas with scores higher than empirically-derived null distributions [14,16,17]. Consistency of voxel activation across varying studies can be assessed due to the fact that ALE values increase with the number of studies reporting activated peaks at a voxel or in close proximity [3]. The cluster-level inference (family-wise error) and the uncorrected *p-*value used to threshold the ALE image were both set to 0.001 (5,000 permutations) in GingerALE.

### Meta-analytic connectivity modelling

MACM investigates whole brain coactivation patterns corresponding to a region of interest (ROI) across a range of tasks. In contrast to resting state functional connectivity analyses, MACM provides a measure of functional connectivity during a range of task-constrained states [28]. Functional connectivity networks can be extracted by functional covariances, in this case during various task paradigms. These networks exhibit interconnected sets of brain regions that interact to perform specific perceptual, motor, cognitive, and affective functions [29]. We used the BrainMap database to search for studies including healthy subjects that report normal mapping activations that exist within the boundaries of a 3-D spherical ROI, regardless of the associated behavioral condition. Whole brain activation coordinates from these selected studies are then assessed for convergence using the ALE method. MACM then yields a map of significant coactivations that provides a task-free meta-analytic model of the region’s functional interactions throughout the rest of the brain [25]. This approach examines brain region co-activity above chance within a given seed region across a large and diverse set of neuroimaging experiments such as those dealing with deception [18,21]. MACM analyses resulting in ALE maps have been validated with diffusion tensor imaging (DTI) and connectivity atlases (CocoMac) [18] and have been demonstrated to be the meta-analytic equivalent of resting-state functional connectivity maps [30,31].

Coordinates of the seven peak activation clusters were identified through D > H ALE and used as seeds for seven subsequent MACM analyses. Using Mango (Multi-image Analysis GUI) [32], binary NIfTI images of 6 mm spherical radius ROIs were created as masks around each peak coordinate. A standard MNI brain template (*Colin27_T1_seg_MNI*.*nii*) was used to visualize the ROI masks. Separate searches for each identified peak ROI were performed using Sleuth. The criteria for each search were: 1) Activations: Activations only, 2) Context: Normal Mapping, 3) Subject Diagnosis: Normals, and 4) the corresponding 6 mm spherical ROI in MNI space. Studies matching this query were downloaded to Sleuth’s workspace. (See S3 Table for specific functional workspaces for each node.) Coordinates from downloaded experiments matching the criteria were analyzed using GingerALE at minimum volume of 250 mm^3^ and a *p*-value < 0.01.

### Network modelling

Network modelling from MACM analyses was carried out using the approach first outlined in Kotkowski et al. [20]. To summarize this procedure, Mango was used to visualize the uncorrected MACM overlay for each seed coordinate on an MNI template (*Colin27_T1_seg_MNI*.*nii*). The uncorrected estimate of meta-analytic connectivity between each seed region and all other specified nodes was extracted and recorded (see raw values in S1 Fig). A Bonferroni correction was used to correct the *p-*value for multiple comparisons between nodes (*p*-value of 0.05/7 = 0.00714). The corrected *p-*values, representing covariance statistics between nodes (i.e. the seed used in each of the seven MACMs) and projections (i.e. the connectivity from the MACM of seed ROI to the six other ROIs), were used to generate the edges in the meta-analytic connectivity model. Connections between the identified peak regions were mapped as nodes exhibiting one-way, two-way, or no significant connections to each other. If only one edge between two nodes was significant (i.e. a significant connection from MACM of ROI 1 to seed 2), the connection was considered unidirectional. On the other hand, if both edges between two nodes were significant (i.e. a significant connection from MACM of seed 1 to ROI 2 and a significant connection from MACM of seed 2 to ROI 1), the connection was considered bidirectional.

### Paradigm class and behavioral domain analyses

Paradigm class and behavioral domain were also analyzed using the resulting nodes from ALE/MACM and the “Paradigm Analysis” and “Behavioral Analysis” plugins for Mango [32]. Paradigm class is a category in BrainMap classifying what experimental task was used. Behavioral domain is a BrainMap category classifying the mental operations likely to be isolated by a given contrast. Laird et al. [33] found that these two fields provide the most salient information for ascertaining a brain region’s function. These analyses assume that the spatial distribution of activation foci derived from BrainMap’s database for each behavioral sub-domain or paradigm class represents that sub-domain’s (or class’s) true probability distribution function [32]. *Z*-scores are generated for observed-minus-expected values for each behavioral sub-domain or paradigm class. Lancaster et al. [32] state that only *z*-scores greater than or equal to 3.0 are significant (comparable to a *p*-value of 0.05 with Bonferroni correction for multiple comparisons). The identification of paradigm class and behavioral domain associated with nodes aids interpretation of connectivity reported via MACM.

## Results

### ALE results for deceptive versus honest behavior

45 studies, 977 foci and 2,836 subjects were included in the ALE meta-analysis to demonstrate activation associated with deceptive versus honest behavior. The D > H ALE revealed seven significant clusters (Table 2). The nearest grey matter associated with each cluster are the left and right insula (L Ins, R Ins), left superior frontal gyrus (L SFG), left and right supramarginal gyrus (L SMG, R SMG), and left and right medial frontal gyrus (L MFG, R MFG). Fig 2 depicts activation of each of the 7 clusters.

**Table 2.**
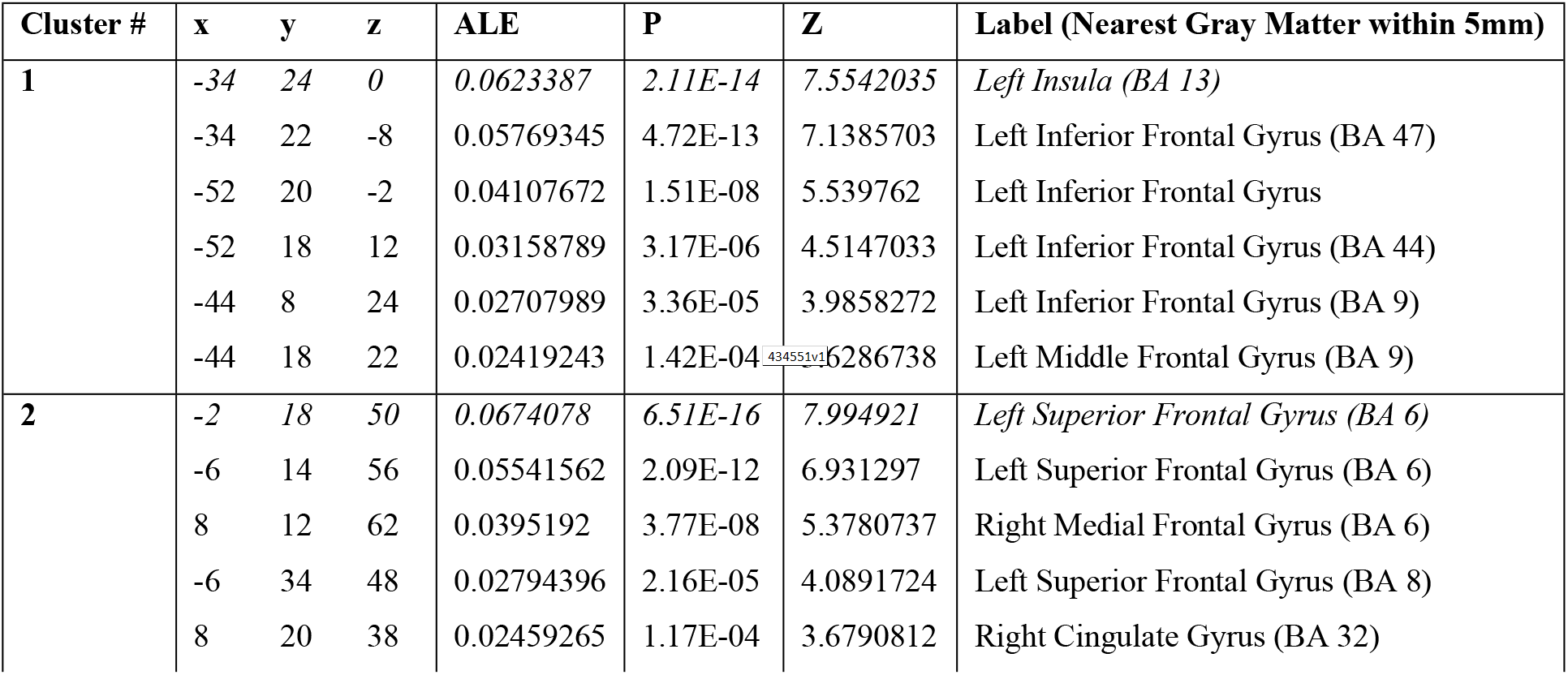
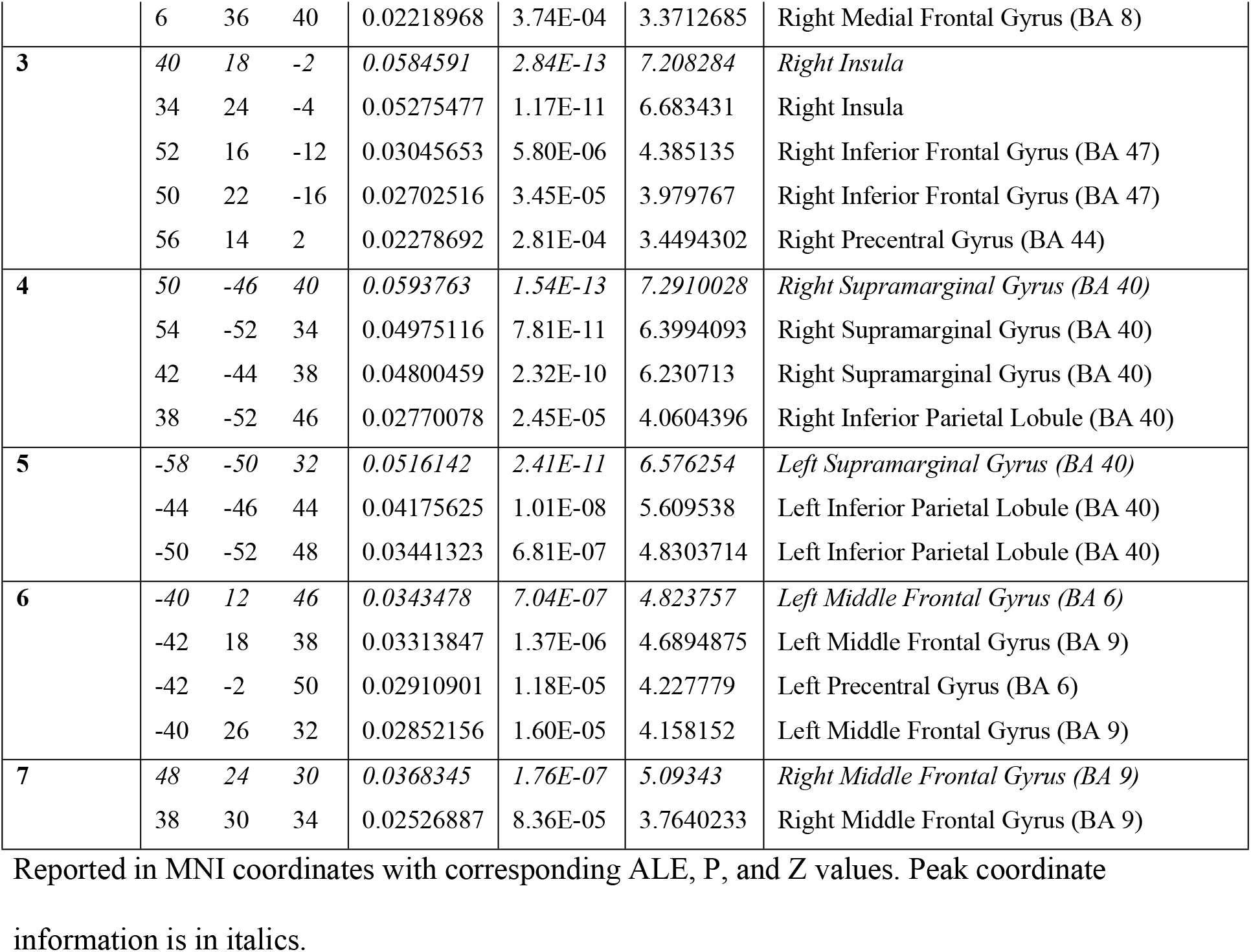
Deceptive > Honest ALE results.

**Fig 2.**
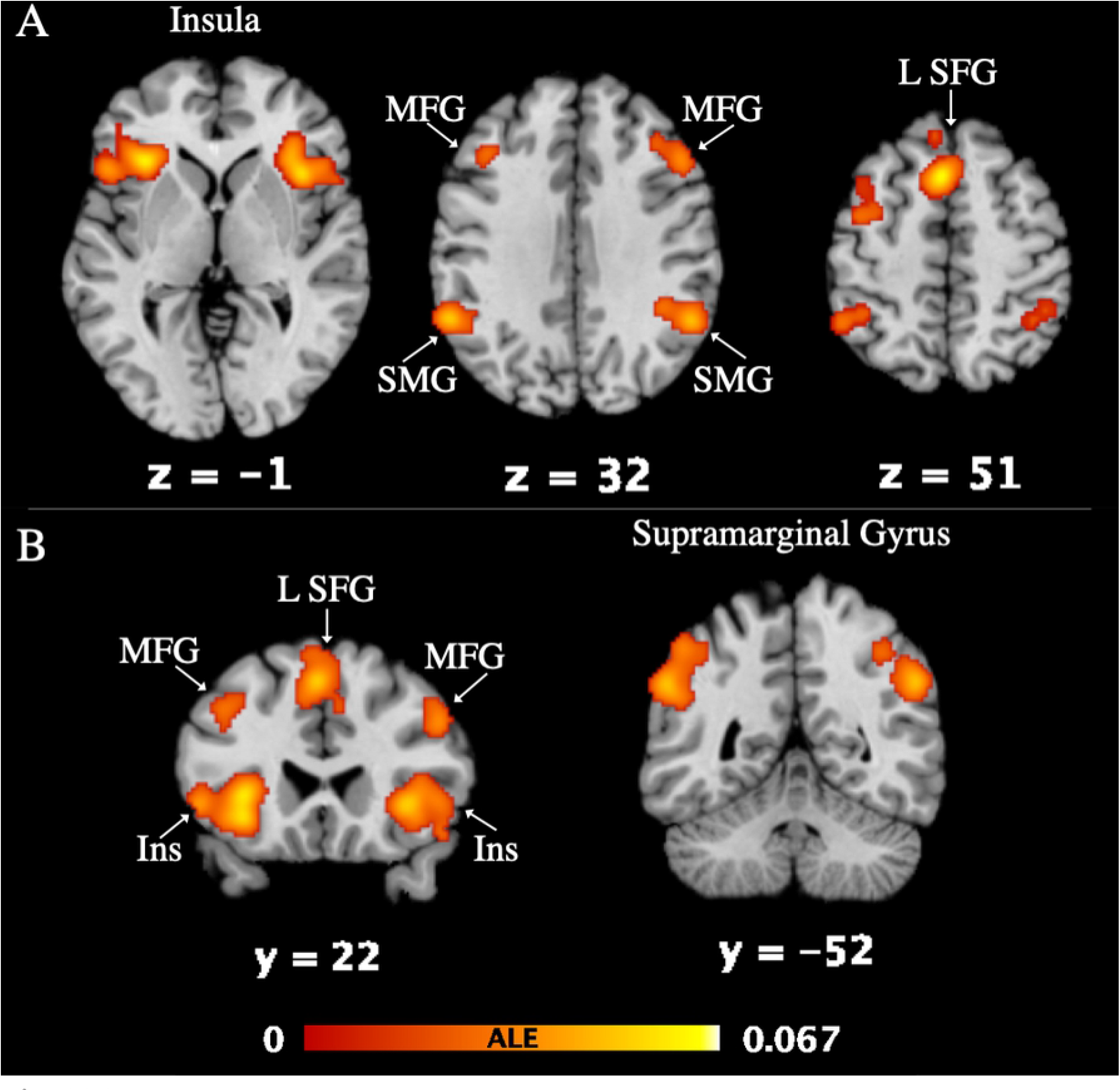
Deceptive > Honest ALE results. Activation is visualized in Mango on a standard MNI brain template (**A**: horizontal slice, **B**: coronal slice; FWE < 0.001, p < 0.001, at 5,000 permutations). Z and Y values correspond to the brain slice label. The activation color (red-yellow) corresponds to the ALE value listed in **Table 2**. Left and right are accurately depicted.

### MACM results for deceptive versus honest behavior

MACM was used to examine the extent of connectivity between the seven clusters identified in the ALE exhibiting greater activation during deception than honest behavior. A unique MACM was carried out for each individual ROI, resulting with seven independent seed to voxel connectivity maps. Bolded lines (Figs 3A and B) represent bidirectionality, indicating that the variance in two nodes is predictive of each other. Arrows (Figs 3A and B) represent unidirectionality, indicating that variance in one node is predictive of variance in another, but not vice versa. The matrix results are shown in Fig 3C (raw scores: S1 Fig).

**Fig 3.**
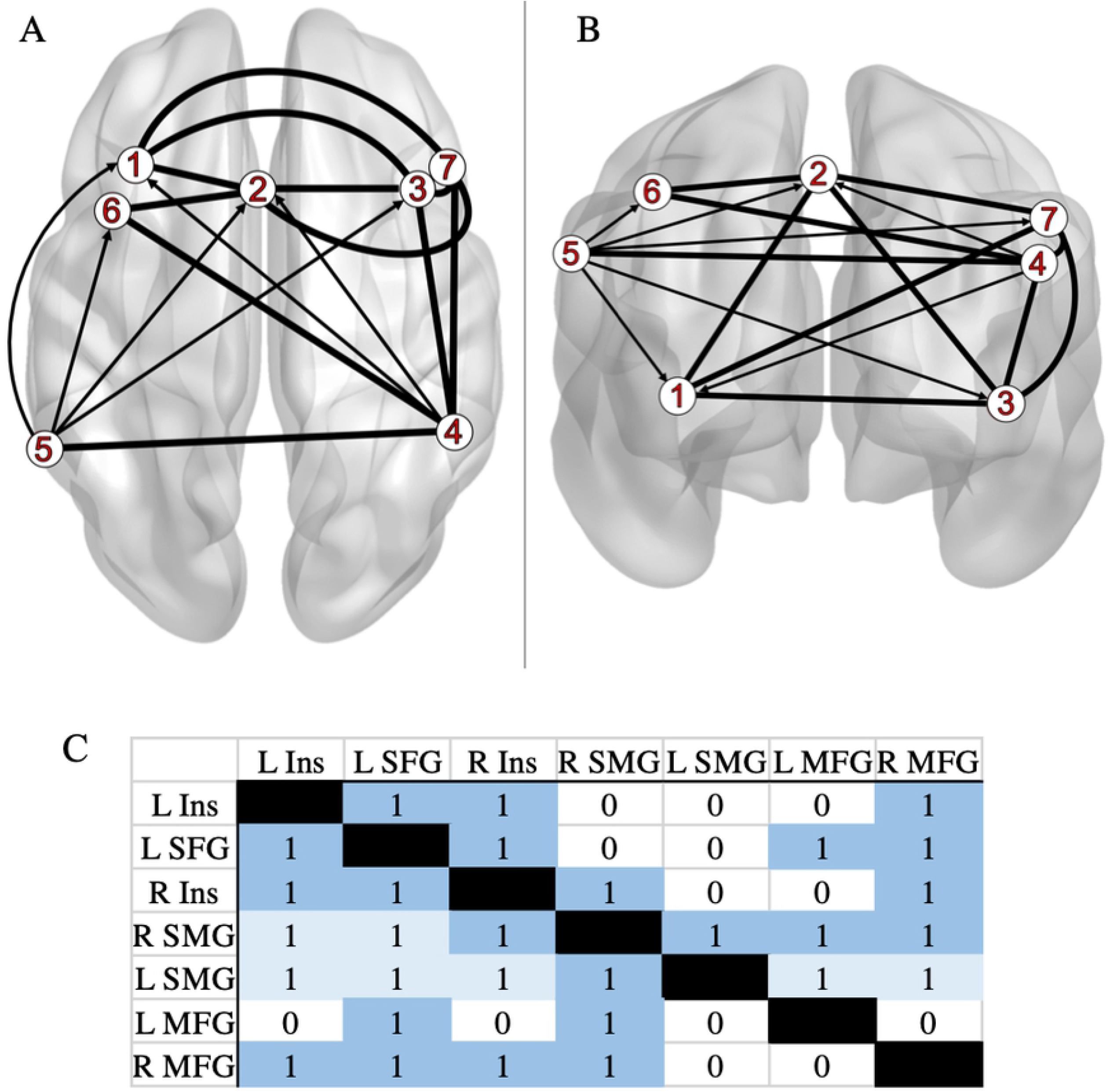
Meta-analytic model of connectivity between Deceptive > Honest peak regions. **A**: horizontal slice and **B**: coronal slice. Data were visualized with the BrainNet Viewer [34] (*http://www.nitrc.org/projects/bnv/*). **Key** (ROI Labels): 1: left insula (L Ins); 2: left superior frontal gyrus (L SFG); 3: right insula (R Ins); 4: right supramarginal gyrus (R SMG); 5: left supramarginal gyrus (L SMG); 6: left medial frontal gyrus (L MFG); 7: right medial frontal gyrus (R MFG). (**C**) The matrix depicting connectivity from seed regions (left column) to the whole brain (“1” dark blue: bidirectional; “1” light blue: unidirectional; “0”: no direction implied).

Significant one-way functional connectivity is shown projecting from: R SMG to L Ins, L SFG, and from L SMG to L Ins, R Ins, L SFG, L MFG, R MFG. Significant two-way functional connectivity is shown involving: L Ins to L SFG, R Ins, R MFG; L SFG to R Ins, L MFG, R MFG; R Ins to R SMG, R MFG; R SMG to L SMG, L MFG, R MFG.

### Paradigm class and behavioral domain results

Using Lancaster et al.’s [32] “Paradigm Class” Mango plugin for analysis of BrainMap’s functional database of healthy subjects, 14 significant paradigm classes were related to the seven nodes identified in the D > H ALE meta-analysis. Fig 4A indicates paradigm classes for which the observed regional number of experiments was higher than expected (compared with the distribution across the BrainMap database). All paradigm classes at a *z*-score of >=2.0 are reported in S4 Table. The left insula has the strongest association with the paradigm class “Reward” (*z* = 4.564). The left SFG has the strongest association with the paradigm class of “Finger Tapping/Button Press” (*z* = 4.905). The right insula has the highest association with the paradigm class “Pain Monitor/Discrimination” (*z* = 5.550). These paradigm class analysis results indicate significant associations of the left and right insula with reward paradigms, in addition to significant associations of left SFG and right insula to semantic discrimination and pain discrimination, respectively.

**Fig 4.**
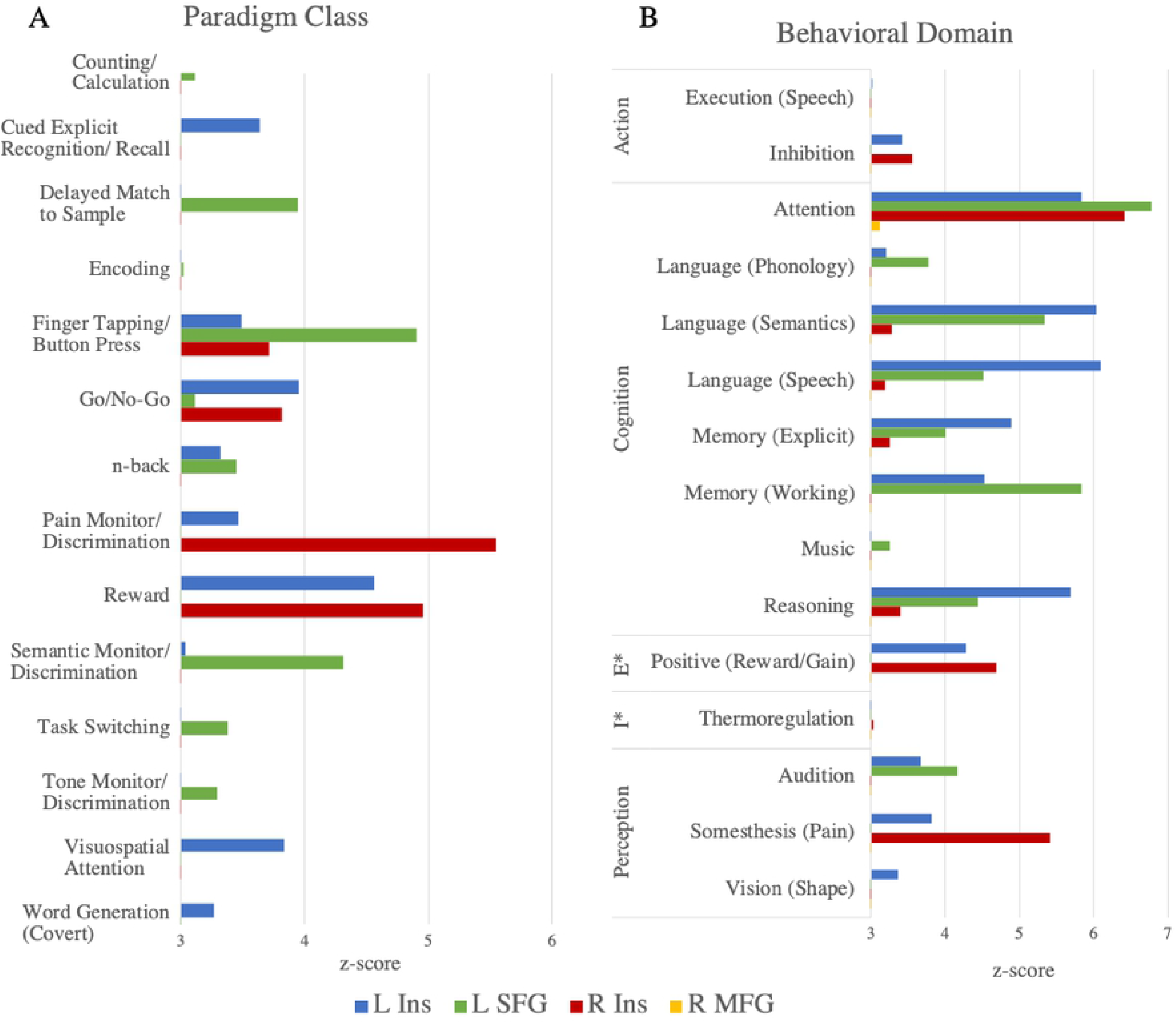
*Z*-scores of (A) paradigm class or (B) behavioral domain analyses. In the Behavioral Domain panel (**B**), the Emotion and Interoception domains are abbreviated as “E” and “I” respectively. Only paradigm classes or behavioral sub-domains passing the threshold of *z* >= 3.0 are depicted.

Subsequent behavioral domain analysis of the seven nodes from ALE/MACM with the “Behavioral Analysis” Mango plugin [32] identified 15 significant sub-domains. Fig 4B indicates behavioral sub-domains (within one of five domains) for which the observed regional number of experiments was higher than expected (compared with the distribution across the BrainMap database). All sub-domains at a *z*-score of >=2.0 are reported in S5 Table. The left insula has the strongest association with sub-domains of “Cognition”, including “Language (Speech)” (*z* = 6.097), “Language (Semantics)” (*z* = 6.037), “Attention” (*z* = 5.837), and “Reasoning” (*z* = 5.693). The left SFG also has strong associations with sub-domains of “Cognition”, including “Attention” (*z* = 6.78), “Memory (Working)” (*z* = 5.829), and “Language (Semantics)” (*z* = 5.335). The right insula has strongest associations with “Attention” of the “Cognition” domain (*z* = 6.421) and “Somesthesis (Pain)” of the “Perception” domain (*z* = 5.417). The right MFG has one significant association with the “Attention” sub-domain of “Cognition” (z = 3.124). These results indicate that the bilateral insula, L SFG, and R MFG are mainly associated with behaviors regarding cognition.

## Discussion

In the presented series of meta-analyses, we conducted activation likelihood estimation and meta-analytic connectivity modelling in addition to subsequent paradigm class and behavioral domain analyses using reported neuroimaging findings for deception tasks.

### Regions associated with deception

The findings of this study align well with previously reported findings while presenting new information regarding functional connectivity of deception-related brain regions. Results from the ALE identified seven brain regions significantly activated during deception, including bilateral insula, left superior frontal gyrus, bilateral supramarginal gyrus, and bilateral medial frontal gyrus. These regions match regions reported in previous meta-analyses: BA 6 (SFG), BA 40 (IPL or SMG), BA 6 (MFG). Our first hypothesis was supported in that the study replicates findings of prefrontal (BA 9 and 13) and memory-related (BA 6) regional activation during deception. Various additional regions were consistently active, most likely resulting from the variety of paradigms included in ALE. Interestingly, the regions that we found to be significantly active during deception tasks matched those reported in the most recent meta-analysis [13]. Here we discuss each region’s functional significance, relationship to sociocognitive behaviors of deception, and make comparisons to existing deception literature.

### Insula

Recent studies using ecologically valid paradigms involved more of the participants’ emotions as evidenced by consistent activation in the insula and other emotion-related brain regions [35]. These recent studies have added evidence that the insula is part of a reflexive, automatic system of social cognition. In Baumgartner et al.’s study [35], results demonstrated increased activation of the anterior insula in dishonest subjects compared to honest subjects. Further, the researchers state that subjects in the dishonest group who later intended to break promises demonstrate increased bilateral frontoinsular cortex activation during that (promise) stage. Proposed reasons for insular activity in dishonesty or deception include insular activation during aversive emotional experiences associated with unfairness, threat of punishment, and anticipation of negative/unknown emotional events [35]. The researchers also state that aversive experiences may include “guilty conscience” towards the other individual who will eventually be misled.

### Superior frontal gyrus

The SFG has been associated with cognitive processes such as working memory, response inhibition, task switching, visual attention, and theory of mind [13]. More specific to deception behavior, Chen et al. [36] reported overlapping SFG activation between feigned short-term and long-term memory. This finding supports the role of SFG in executive function aspects of feigned memory impairment, whether short-term or long-term memory [36]. In addition, Yin et al. [37] reported that both spontaneous and instructed lying coactivate the SFG among other regions. Researchers also report the involvement of SFG in identity faking aspects of deception behavior [38]. Since SFG has implications with working memory, Ding et al. [38] state that both SFG and working memory functions play a role in deceptively faking one’s identity.

### Supramarginal gyrus

The supramarginal gyrus lies within the inferior parietal lobule, an area commonly associated with deception since the pioneering neuroimaging study by Spence et al. [6]. Instructed deception has been shown to involve the IPL [37]. Various other studies have associated the inferior parietal regions with the execution of deception. Ito et al. [39] reported increased SMG activity in the execution phase of a deception task compared with telling the truth. Kireev et al. [40] found a similar result in that a network including the IPL demonstrated increased activation during deliberate deception processing/execution. In addition, Ofen et al. [41] found similar activation of parietal regions during the execution of a deceptive response. Potential reasons for the involvement of SMG/IPL in executing deception include parietal regions supporting executive functioning (i.e. working memory) [39] and cognitive control processes as they are commonly activated during tasks that require high levels of cognitive control [41]. Further evidence of this comes from a study where activation of parietal regions was associated with intentional feigned responses and not unintentional errors [41].

It has also been suggested that SMG/IPL is engaged when detecting salient stimuli and processing judgements regarding deception [10] as well as probability monitoring and response counting [5]. Browndyke et al. [5] state that these sociocognitive aspects may allow the deceiver to lie less obviously, or better feign an impairment. Further, the study participants subsequently reported attempts to gauge the proportion of their true versus feigned responses in order to create less detectable deception [5]. Along this line of thought, the parietal regions (SMG/IPL) have been associated with theory of mind [13]. Theory of mind necessitates the ability to understand and predict another individual’s behavior (via inferences regarding mental state, intentions, feelings, expectations, beliefs, or knowledge) and to cognitively represent one’s own mental state [42]. Evidence of the association between SMG and the sociocognitive process of theory of mind includes the activation of SMG in pro-social lying that was deemed morally appropriate [43] and the recruitment of IPL regions for top-down modulation of emotional responses [44].

### Medial frontal gyrus

Frontal (namely prefrontal) regions have markedly been reported in association with deception tasks and behaviors. Sun et al. [45] demonstrated that lies elicited stronger MFG activation compared to truth. Moreover, Bhatt et al. [46] state that MFG may play a role in familiarity-based deception (rather than familiarity or deception individually). Liu et al. [47] stated that (left) MFG seemed to be primarily responsible for the falsification process in conditional proposition testing. The researchers noted the association between MFG and working memory and higher-level control processes (i.e. coordinating widely distributed cognitive and emotional reactions, learning new rules, and processing logical relationships) [47]. Further, involvement of frontal lobe regions is consistent with the conceptualization of deception as an executive control incentive task [11,48].

### Connectivity analyses

Our second hypothesis was also supported by the involvement of the prefrontal and memory-related regions in the connectivity model. The connectivity modelling used in the current meta-analysis, which adds new information regarding deception-related brain regions, has not been done in this realm of research before to our knowledge. MACM of brain regions active during deception, identified via ALE, show that these regions are also highly connected to each other. Each of the seven nodes were involved in at least one significant bidirectional connection. Interestingly, only the seed nodes for left and right supramarginal gyri projected to other nodes (in other words, were involved in unidirectional connections). Thus, activation of SMG is likely predictive of activation in bilateral insula, left SFG, or bilateral MFG (respectively). This means that the bilateral SMG must engage with other regions to engage in deception tasks, however those other regions are not required for deception. Other regions identified in our deception ALE (i.e. bilateral insula, left SFG, bilateral MFG) likely have supportive roles in cognitive aspects of the tasks. This may well be the case since, in order to lie, an individual must construct new information while withholding factual information during a social interaction with another individual [49]. The important role SMG plays in deception is further supported by our paradigm class and behavioral domain findings. The bilateral SMG did not elicit significant (*z*-score >= 3.0) paradigm class or behavioral domain information that would indicate SMG involvement in other cognitive/task-based aspects in the current meta-analysis. Together, the connectivity model, paradigm class, and behavioral domain findings of the current study could implicate the supramarginal gyrus as a key region in a brain network that allows individuals to successfully deceive one another.

### Importance of neuroimaging deception and its application

A major motivation behind the study of deception is the ability to reliably detect when a given individual is being truthful or is lying [11]. The law often concerns itself with this phenomenon as it contributes to judgements regarding human behavior. Untruthful statements are possible and commonly made by plaintiffs, defendants, and witnesses alike [50]. Assessing the veracity of statements made by individuals inside and outside of the courtroom is a crucial component of just and efficient legal resolution [50]. Legal actors increasingly offer neuroscientific evidence during litigation and policy discussions. Similarly, cognitive neuroscientists aim to address important problems confronted by the law by explaining neuropsychological mechanisms that give rise to thoughts and actions [51]. The utility of neuroscientific evidence depends both on the accuracy of the neuroscience as well as the appropriate usage by legal actors. Though specific courtroom scenarios deal with individuals, group-level studies are needed as fMRI-based evidence will be used to establish the reliability of instances related to any deception apparent in court [50]. Accurate detection of deception in humans is of particular importance in ensuring valid and just forensic practices and legal proceedings.

Where the legal system and neuroscience overlap is in the attempts to utilize neuroscientific advances to yield better answers to legally relevant questions that have had historically unsatisfying solutions [51]. Some questions include whether or not an individual is responsible for their behavior, if an individual is competent, what an individual remembers, and pertaining to the current meta-analysis, if an individual is lying. Legal cases from the last decade or so have involved methods of brain-based lie detection, brain-based memory detection (wherein under controlled experimental conditions memory states may be detected using fMRI data), detection and classification of “culpable mental states” including purposeful, knowing, reckless, and negligent (based on the “Model Penal Code”), and investigations of the decision-making processes of, not only if an individual is criminally liable, but also how to then punish that individual in an unbiased and just fashion [51]. However, all of these aspects pertaining to criminal law have their apparent downfalls (for more on this see [51]). Those at the intersection of neuroscience and the law (commonly called “neurolaw”) focus on non-criminal law as well: the aging brain in regard to wills, trusts, and estates; disability and social security laws in association with the neuroscience of pain; similarly, brain injury cases and medical malpractice; and more.

Neuroimaging has been used in legal proceedings since the early twentieth century, with use of electroencephalography (EEG) appearing in the 1940s, computed tomography (CT) appearing in 1981, positron emission tomography (PET) appearing in 1992, and fMRI not long after [52]. Over the last two decades alone, the use of neuroscientific evidence in general and neuroimaging-based evidence specifically has increased tremendously in the United States [52]. Jones [53] has identified seven categories for the applications of neuroscience to the legal setting: buttressing, detecting, sorting, challenging, intervening, explaining, and predicting. We believe this meta-analytic view of deception fits into the detecting and explaining categories, wherein neuroscience is used to gain otherwise elusive insights and to shed light on not well understood phenomenon. Our work contributes to efforts of detecting deception-based activity in the functional brain rather than the activity of the nervous system (i.e. heart rate/blood pressure, respiration, skin conductivity, etc. used in polygraphy). Benefits of this have been reviewed at length [54]. In agreement with what is written in a recent review [51], we believe that there is a common ground where the long-term effects of neuroscience on law are not overstated but we can appropriately consider that neuroscience has something useful to offer the legal system.

### Challenges and limitations

Spence et al. [49] predicted the problems that have persisted in the neuroimaging literature of deception: 1) ecological validity: the experiments generally involve compliant subjects who are not involved in high-stakes situations that pertain to forensics or the legal system (thus, these studies are unable to address how the brain functions when someone is intentionally lying to cause harm or deceive for a known purpose and may not extrapolate to circumstances wherein deception is an automatic process driving malevolent behavior) [40]; 2) experimental design: some experiments have simple designs of simulated deception that facilitate simple contrasts (lie > truth) which may not cohere in the real world (where there exists imprecise information, mixed motives, etc.); 3) statistical power: there may well be a range of individual differences that would make it premature to extrapolate from neuroimaging data to an individual suspect in a courtroom.

The current meta-analysis regarding brain regions active during deceptive versus honest behavior addresses the above problems to some degree by including ecologically valid studies in our total pool and drawing results from a large, heterogenous sample. More recently, studies and their respective paradigms have attempted to evoke “realistic social exchanges” by allowing participants the free choice to break or keep a promise, mitigating to some degree the previous work. These ecologically valid studies were included in our current meta-analyses. Also, the nature of coordinate-based meta-analyses that include task-based studies allows results to be drawn from a large, heterogeneous sample. This takes into account paradigms that may or may not involve compliant subjects in somewhat realistic circumstances, and that may or may not include “simple contrasts”, as long as the inclusion criteria are met. Regarding statistical power, the recommended number of included experiments has been met in the current meta-analysis (20 experiments in order to achieve sufficient statistical power) [55].

### Future Directions

Due to the previously noted association of supramarginal gyrus and theory of mind aspects of deception, a potential next step could be analyzing regions found in the current study with regions involved in theory of mind. Deception is related to theory of mind, as deceiving another individual necessitates knowledge of the victims’ thoughts and beliefs as well as analysis of responses to the lie made in the social context [11]. Thus, follow-up meta-analyses can be conducted and subsequently compared to the findings of the current study to determine if overlapping regional activation exists. Of particular interest in such a comparison would be the SMG and SFG which have been associated with theory of mind aspects of deception.

## Conclusion

The current study utilized activation likelihood estimation and the novel approaches of meta-analytic connectivity analysis, paradigm class analysis, and behavioral domain analysis to investigate neuroanatomical correlates of deception and their functional connectivity. Across the varying studies involving differences in context of deception, motivation for deception, response modality, and more, we found significant activation in the insula, superior and medial frontal gyri, and supramarginal gyrus. Moreover, the connectivity model and paradigm/behavioral analyses demonstrate the key role that the supramarginal gyrus has in the brain network associated with deceptive acts and behaviors. An understanding of the neurobiological aspects of deception has implications for subsequent theory of mind and social cognition research in addition to forensic/legal analyses of guilt and responsibility.

## Acknowledgements

We acknowledge and are grateful for the help of Noah Waller, Laura Ireland, Ariana White, and Jenelle Delfino in coding papers for Sleuth, instruction for analyses, and preparing the manuscript. This work is supported by Fund IGF062 Program Neuroscience and Behavioral Health at the University of New Hampshire.

---

**S1 Fig. Raw values**. The uncorrected estimate of meta-analytic connectivity between each seed region and all other specified nodes.

**S1 Table. Contrasts included in ALE of all contrasts (D > H, H > D, etc.)**. This ALE consisted of 46 studies and 202 experiments with 1,423 foci from 4,678 participants.

**S2 Table. Results from ALE of all contrasts**. Contrasts included Deceptive > Honest, Honest > Deceptive, etc.

**S3 Table. Functional MACM workspace information for each node**.

**S4 Table. Paradigm class analysis results**. Using “Paradigm Analysis” plugin for Mango [32]. *Z*-scores that are significant according to Lancaster et al. [32], meaning they have a *z*-score of >= 3.0, are in bold. However, all *z*-scores >= 2.0 are reported here.

**S5 Table. Behavioral domain analysis results**. Using “Behavioral Analysis” plugin for Mango [32]. *Z*-scores that are significant according to Lancaster et al. [32], meaning they have a *z*-score of >= 3.0, are in bold. However, all *z*-scores >= 2.0 are reported here.

**S6 Table. PRISMA Checklist**.

## References

1. Merriam-Webster.com [Internet]. Deception. Merriam-Webster dictionary. [cited 2021 Jan 11]. Available from https://www.merriam-webster.com/dictionary/deception

2. Zuckerman M, Koestner R, Driver R. Beliefs about cues associated with deception. J Nonverbal Behav. 1981 Dec;6(2):105–114. https://doi.org/10.1007/BF00987286.

3. Abe N. How the brain shapes deception: an integrated review of the literature. Neuroscientist. 2011 Oct;17(5):560–74. https://doi.org/10.1177/1073858410393359

4. Hare RD, Neumann CS. Psychopathy as a clinical and empirical construct. Annu Rec Clin Psychol. 2008;4:217–46. https://doi.org/10.1146/annurev.clinpsy.3.022806.091452

5. Browndyke JN, Paskavitz J, Sweet LH, Cohen RA, Tuker KA, Welsh-Bohmer KA, et al. Neuroanatomical correlates of malingered memory impairment: event-related fMRI of deception on recognition memory task. Brain Inj. 2008 Jun;22(6):481–9. https://doi.org/10.1080/02699050802084894

6. Spence SA, Farrow TF, Herford AE, Wilkinson ID, Zheng Y, Woodruff PW. Behavioural and functional anatomical correlates of deception in humans. Neuroreport. 2001 Sep 17;12(13):2849–53. https://doi.org/10.1097/00001756-200109170-00019

7. Langleben DD, Schroeder L, Maldjian JA, Gur RC, McDonald S, Ragland JD, et al. Brain activity during simulated deception: an event-related functional magnetic resonance study. Neuroimage. 2002 Mar;15(3):727–32. https://doi.org/10.1006/nimg.2001.1003

8. Lee TMC, Liu HL, Tan LH, Chan CC, Mahankali S, Feng C, et al. Lie detection by functional magnetic resonance imaging. Hum Brain Mapp. 2002 Mar;15(3):157–64. https://doi.org/10.1002/hbm.10020

9. Ganis G, Kosslyn SM, Stose S, Thompson WL, Yurgelun-Todd DA. Neural correlates of different types of deception: an fRMI investigation. Cereb Cortex. 2003 Aug;13(8):830–6. https://doi.org/10.1093/cercor/13.8.830

10. Cui Q, Vanman EJ, Wei D, Yang W, Jia L, Zhang Q. Detection of deception based on fMRI activation patterns underlying the production of a deceptive response and receiving feedback about the success of the deception after a mock murder crime. Soc Cogn Affect Neurosci. 2014 Oct;9(10):1472–80. https://doi.org/10.1093/scan/nst134

11. Christ SE, Van Essen DC, Watson JM, Brubaker LE, McDermott KB. The contributions of prefrontal cortex and executive control to deception: evidence from activation likelihood estimate meta-analyses. Cereb Cortex. 2009 Jul;19(7):1557–66. https://doi.org/10.1093/cercor/bhn189

12. Lisofsky N, Kazzer P, Heerkeren HR, Prehn. Investigating socio-cognitive processes in deception: a quantitative meta-analysis of neuroimaging studies. Neuropsychologia. 2014 Aug;61:113–22. https://doi.org/10.1016/j.neuropsychologia.2014.06.001

13. Yu J, Tao Q, Zhang R, Chan CCH, Lee TMC. Can fMRI discriminate between deception and false memory? A meta-analytic comparison between deception and false memory studies. Neurosci Biobehav Rev. 2019 Sep;104:43–55. https://doi.org/10.1016/j.neubiorev.2019.06.027

14. Eickhoff SB, Bzdok D, Laird AR, Kurth F, Fox PT. Activation likelihood estimation meta-analysis revisited. Neuroimage. 2012 Feb 1;59(3):2349–61. https://doi.org/10.1016/j.neuroimage.2011.09.017.

15. Laird AR, Eickhoff SB, Kurth F, Fox PM, Uecker AM, Turner JA, et al. ALE meta-analysis workflows via the BrainMap database: Progress towards a probabilistic functional brain atlas. Front Neuroinform. 2009 Jul 9;3:23. https://doi.org/10.3389/neuro.11.023.2009

16. Laird AR, Fox PM, Price CJ, Glahn DC, Uecker AM, Lancaster JL et al.ALE meta-analysis: controlling the false discovery rate and performing statistical contrasts. Hum Brain Mapp. 2005 May;25(1):155–64. https://doi.org/10.1002/hbm.20136

17. Turkeltaub PE, Eickhoff SB, Laird AR, Fox M, Wiener M, Fox P. Minimizing within– experiment and within-group effects in Activation Likelihood Estimate meta-analyses. Hum Brain Mapp. 2012 Jan;33(1):1–13. https://doi.org/10.1016/j.physbeh.2017.03.040

18. Robinson JL, Laird AR, Glahn DC, Lovallo WR, Fox PT. Meta-analytic connectivity modelling: Delineating the functional connectivity of the human amygdala. Hum Brain Mapp. 2009 Jan 14;31(2):173–84. https://doi.org/10.1002/hbm.20854

19. Robinson JL, Laird AR, Glahn DC, Blangero J, Sanghera MK, Pessoa L, et al. The functional connectivity of human caudate: an application of meta-analytic connectivity modeling with behavioral filtering. Neuroimage. 2012 Mar;60(1):117–29. https://doi.org/10.1016/j.neuroimage.2011.12.010

20. Kotkowski E, Price LR, Fox LR, Vanasse TJ, Fox PT. The hippocampal network model: A transdiagnostic metaconnectomic approach. Neuroimage Clin. 2018 Jan 8;18:115–129. https://doi.org/10.1016/j.nicl.2018.01.002

21. Eickhoff SB, Jbabdi S, Caspers S, Laird AR, Fox PT, Zilles K, et al. Anatomical and functional connectivity of cytoarchitectonic areas within the human parietal operculum. J Neurosci. 2010 May 5;30(18):6409–21. https://doi.org/10.1523/JNEUROSCI.5664-09.2010

22. Moher D. Liberati A, Tetzlaff J, Altman DG, PRISMA Group. Preferred reporting items for systematic reviews and meta-analyses: the PRISMA statement. PLoS Med. 2009 Jul 21;6(7):e1000097. https://doi.org/10.1371/journal.pmed1000097

23. Eickhoff SB, Laird AR, Grefkes C, Wang LE, Zilles K, Fox PT. Coordinate-based activation likelihood estimation meta-analysis of neuroimaging data: A random-effects approach based on empirical estimates of spatial uncertainty. Hum Brain Mapp. 2009 Jan;30(9):2907–26. https://doi.org/10.1002/hbm.20718

24. Fox PT, Lancaster JL. Mapping context and content: the BrainMap model. Nat Rev Neurosci. 2002 Apr;3(4):319–21. https://doi.org/10.1038/nrn789

25. Laird AR, Eickhoff SB, Fox PM, Uecker AM, Ray KL, Saenz JJ. The BrainMap strategy for standardization, sharing, and meta-analysis of neuroimaging data. BMC Res Notes. 2011 Sep 9;4:349. https://doi.org/10.1186/1756-0500-4-349

26. Fox PT, Laird AR, Fox SP, Fox PM, Uecker AM, Crank M, et al. BrainMap taxonomy of experimental design: description and evaluation. Hum Brain Mapp. 2005 May;25(1):185–98. https://doi.org/10.1002/hbm.20141

27. Vanasse TJ, Fox PM, Barron DS, Robertson M, Eickhoff SB, Lancaster JL, et al. BrainMap VBM: an environment for structural meta-analysis. Hum Brain Mapp. 2018 Aug;39(8):3308–3325. https://doi.org/10.1002hbm.24078

28. Langer R, Rottschy C, Laird AR, Fox PT, Eickhoff SB. Meta-analytic connectivity modeling revisited: controlling for activation base rates. Neuroimage. 2014 Oct 1;99:559–70. https://doi.org/10.1016/j.neuroimage.2014.06.007

29. Bressler SL, Menon V. Large-scale brain networks in cognition: emerging methods and principles. Trends Cogn Sci. 2010 Jun;14(6):277–90. https://.doi.org/10.1016/j.tics.2010.04.004

30. Laird AR, Eickhoff SB, Karl L, Robin DA, Glahn DC, Fox PT. Investigating the functional heterogeneity of the default mode network using coordinate-based-meta-analytic modeling. J Neurosci. 2009 Nov 18;29(46):14496–505. https://doi.org/10.1523/JNEUROSCI.4004-09.2009

31. Smith SM, Fox PT, Miller KL, Glahn DC, Fox PM, Mackay CE, et al. The functional architecture of the human brain: correspondence between resting fMRI and task-activation studies. Proc Natl Acad Sci U S A. 2009 Aug 4;106(31):13040–45. https://doi.org/10.1073/pnas.0905267106

32. Lancaster JL, Laird AR, Eickhoff SB, Martinez MJ, Fox PM, Fox PT. Automated regional behavioral analysis for human brain images. Front Neuroinform. 2012 Aug 28;6:23. https://doi.org/10.3389/fninf.2012.00023

33. Laird AR, Fox PM, Eickhoff SB, Turner JA, Ray KL, McKay DR, et al. Behavioral interpretations of intrinsic connectivity networks. J Cogn Neurosci. 2011 Dec;23(12):4022–37. https://doi.org/10.1162/jocn_a_00077

34. Xia M, Wang J, He Y. BrainNet Viewer: a network visualization tool for human brain connectomics. PLoS One. 2013 Jul 4;8(7):e68910. https://doi.org/10.1371/journal.pone.0068910

35. Baumgarter T, Fischbacher U, Feirabend A, Lutz K, Fehr R. The neural circuitry of a broken promise. Neuron. 2009 Dec 10;64(5):756–70. https://doi.org/10.1016/j.neuron.2009.11.017

36. Chen Z-X, Xue L, Liang X-Y, Mei W, Zhang Q, Zhao H. Specific marker of feigned memory impairment: The activation of left superior frontal gyrus. J Forensic Leg Med. 2015 Nov;36:164–71. https://doi.org/10.1016/j.jflm.2015.09.008

37. Yin L, Reuter M, Weber B. Let the man choose what to do: Neural correlates of spontaneous lying and truth-telling. Brain Cogn. 2016 Feb;102:13–25. https://doi.org/10.1016/j.bandc.2015.11.007

38. Ding XP, D. X, Lei D, Hu CS, Fu G, Chen G. The neural correlates of identity faking and concealment: an fMRI study. PLoS One. 2012;7(11):e48639. https://doi.org/10.1371/journal.pone.0048639

39. Ito A, Abe N, Fujii T, Hayashi A, Ueno A, Mugikura A, et al. The contribution of the dorsolateral prefrontal cortex to the preparation for deception and truth-telling. Brain Res. 2012 Jun 29;1464:43–52. https://doi.org/10.1016/j.brainres.2012.05.004

40. Kireev M, Korotkov A, Medvedeva N, Medvedev S. Possible role of an error detection mechanism in brain processing of deception: PET-fMRI study. Int J Psychophysiol. 2013 Dec;90(3):291–9. https://doi.org/10.1016/j.ijpsycho.2013.09.005

41. Ofen N, Whitfield-Gabrieli S, Chai XJ, Schwarlose RF, Gabrieli JDE. Neural correlates of deception: lying about past event and personal beliefs. Soc Cogn Affect Neurosci. 2017 Jan 1;12(1):116–127. https://doi.org/10.1093/scan/nsw151

42. Lissek S, Peters S, Fuchs N, Witthaus H, Nicolas V, Tegenthoff M, et al. Cooperation and deception recruit different subsets of the theory-of-mind network. PLoS One. 2008 Apr 23;3(4):e2023. https://doi.org/10.1371/journal.pone.0002023

43. Hayashi A, Abe N, Fujii T, Ito A, Ueno A, Koseki Y, et al. Dissociable neural systems for moral judgment of anti-and pro-social lying. Brain Res. 2014 Mar 27;1556:46–56. https://doi.org/10.1016/j.brainres.2014.02.011

44. Lee TMC, Lee TMY, Raine A, Chan CCH. Lying about the valence of affective pictures: an fMRI study. PLoS One. 2010 Aug 25;5(8):e12291. https://doi.org/10.1371/journal.pone.0012291

45. Sun D, Lee TMC, Chan CCH. Unfolding the spatial and temporal neural processing of lying about face familiarity. Cereb Cortex. 2015 Apr;25(4):927–v 36. https://doi.org/10.1093/cercor/bht284

46. Bhatt S, Mbwana J, Adeyemi A, Sawyer A, Hailu A, VanMeter J. Lying about facial recognition: an fMRI study. Brain Cogn. 2009. Mar;69(2):382–90. https://doi.org/10.1016/j.bandc.2008.08.033

47. Liu J, Zhang M, Jou J, Wu X, Li W, Qiu J. Neural bases of falsification in conditional proposition testing: evidence from an fMRI study. Int J Psychophysiol. 2012 Aug;85(2):249–56. https://doi.org/10.1016/j.ijpsycho.2012.02.011

48. Miyake A, Friedman NP, Emerson MJ, Witzki AH, Howerter A, Wager TD. The unity and diversity of executive functions and their contributions to complex “Frontal Lobe” tasks: a latent variable analysis. Cogn Psychol. 2000 Aug;41(1):49–100. https://doi.org/10.1006/cogp.1999.0734

49. Spence SA, Hunter MD, Farrow TFD, Green RD, Leung DH, Hughes CJ, et al. A cognitive neurobiological account of deception evidence from functional neuroimaging. Philos Trans R Soc Lond B Biol Sci. 2004 Nov 29;359(1451):1755–62. https://doi.org/10.1098/rstb.2004.1555

50. Adelsheim C. Functional magnetic resonance detection of deception: Great as fundamental research, inadequate as substantive evidence. Mercer Law Rev. 2011;62(3):6.

51. Jones OD, Wagner AD. Law and neuroscience: Progress, promise, and pitfalls. In: Gazzaniga MS, Mangum GR, D Poeppel, editors. The cognitive neurosciences. 6th ed. The MIT Press; 2018.

52. Aono D, Yaffe G, Kober H. Neuroscientific evidence in the courtroom: a review. Cogn. Research. 2019;4(40). https://doi.org/10.1186/s41235-019-0179-y

53. Jones OD. Seven ways neuroscience aids law. In Battro AM, Dehaene S, Sorondo MS, Singer WJ, editors. Neurosciences and the human person: new perspectives on human activities. Vatican City: The Pontifical Academy of Sciences; 2013. p. 181–194.

54. Farah MJ, Hutchinson JB, Phelps EA, Wagner AD. Functional MRI-based lie detection: scientific and societal challenges. Nat Rec Neurosci. 2014 Feb;15(2):123–31. https://doi.org/10.1038/nrn3665

55. Eickhoff SB, Nichols TE, Laird AR, Hoffstaedter F, Amunts K, Fox PT, et al. Behavior, sensitivity, and power of activation likelihood estimation characterized by massive empirical simulation. Neuroimage. 2016 Aug 15;137:70–85. http://dx.doi.org/10.1016/j.neuroimage.2016.04.072

